# Behavioral Time Scale Plasticity of Place Fields: Mathematical Analysis

**DOI:** 10.1101/2020.09.11.293787

**Authors:** Harel Z. Shouval, Ian Cone

## Abstract

Traditional synaptic plasticity experiments and models depend on tight temporal correlations between pre- and postsynaptic activity. These tight temporal correlations, on the order of tens of milliseconds, are incompatible with significantly longer behavioral time scales, and as such might not be able to account for plasticity induced by behavior. Indeed, recent findings in hippocampus suggest that rapid, bidirectional synaptic plasticity which modifies place fields in CA1 operates at behavioral time scales. These experimental results suggest that presynaptic activity generates synaptic eligibility traces both for potentiation and depression, which last on the order of seconds. These traces can be converted to changes in synaptic efficacies by the activation of an instructive signal that depends on naturally occurring or experimentally induced plateau potentials. We have developed a simple mathematical model that is consistent with these observations. This model can be fully analyzed to find the fixed points of induced place fields, the convergence to these fixed points, and how these fixed points depend on system parameters such as the size and shape of presynaptic place fields, the animal’s velocity, and the parameters of the plasticity rule.

## 1 Introduction

Experiments and models of synaptic plasticity have, for several decades, concentrated on plasticity which depends on coincident or nearly coincident activation of pre- and postsynaptic cells. This is most clearly exemplified by spike timing dependent plasticity (STDP), in which timing differences between pre- and postsynaptic spikes, on the order of tens of milliseconds, significantly impact the sign and magnitude of synaptic plasticity [1, 2, 3]. Such correlations between pre- and postsynaptic activity are possible biological implementations of unsupervised learning [4, 5]. However, many aspects of behavioral plasticity depend on a supervising signal, or a reward, which can occur with delays that range from hundreds of milliseconds to seconds or more. The difficulty in associating events (such as stimulus and reward) at larger time scales is called the temporal credit assignment problem [6]. Various methods to solve the temporal credit assignment problem have been proposed, none of which solely depend on coincidences on the range of tens of milliseconds. One possible solution depends on synaptic eligibility traces, which can last for several seconds following neural activity, and which can be converted into changes in synaptic efficacies if they are followed by a reward or an instructive signal.

Recent evidence in several systems has provided experimental support for the existence of synaptic eligibility traces. It has been shown that a neuromodulator applied seconds after a pre before post pairing protocol can induce long-term potentiation (LTP), and that this depends on delayed application of neuromodulator [7, 8, 9, 10]. It has also been shown that after a post before pre pairing protocol, a different neuromodulator can induce long-term depression (LTD) [8]. These results indicate that pairing of pre- and post-synaptic activities can generate some currently undetermined biochemical processes, which last for several seconds, that are the substrates of the synaptic eligibility traces for LTP and LTD. If a neuromodulator is applied while the trace is sufficiently active, either LTP or LTD is induced, depending on the details of the pairing protocol. Note that in these experiments, the traces induced depend on both pre- and post-synaptic activity, while the conversion of these traces into efficacy changes depend on a third factor, a neuromodulatory signal. These examples are therefore examples of three-factor learning [11]. Theoretical models consistent with these experimental observations have been shown to be useful in accounting for learning in model networks [12, 8, 13]. In the cerebellum, a structure indicated in some forms of conditioning [14], learning depends on two factors, the activity in the parallel fiber pathway and in the climbing fibers. However, these two factors are not pre and postsynaptic activity, as the climbing fiber activity acts as an instructive signal. These specific roles for the two pathways play a role in conditioning, since parallel fiber activity depends on the conditioned stimulus and climbing fiber activity depends on the unconditioned stimulus or possibly the prediction error [15]. Learning in this system is accomplished through LTD, and the induction of LTD is maximized when parallel fiber activity precedes climbing fiber activity by 50-200ms. The mechanisms for this small delay, which is reminiscent of a short trace, are mediated directly through calcium transients and are well understood [16].

Recent plasticity experiments in hippocampus in vivo [17, 18] have shown place-field plasticity that occurs rapidly in response to either naturally occurring or artificially induced dendritic calcium spikes, also known as ”plateau potentials”. These protocols have shown both an increase and a decrease in synaptic efficacies occurring in synapses that were active seconds before or after the plateau potentials. This plasticity, coined ”behavioral time synaptic plasticity” (BTSP), is therefore also likely to depend on synaptic eligibility traces, both for LTP and LTD. A recent paper has shown that these traces likely depend only on presynaptic activity and the existing synaptic efficacy, and that change in synaptic efficacies depends on the overlap between these traces and an instructive signal that is activated by the plateau potential[18]. The data therefore supports a two-factor model in which the two factors are presynaptic activity and an instructive signal.

The model we present and analyze here stems from these previous experimental and theoretical results. We show that the place fields produced by the model have fixed points, that these fixed points can be defined and calculated, and that the convergence rate to these fixed points can be estimated. In some simple cases these fixed points can be fully solved analytically. Using these solutions, we show how these fixed points depend on the system’s parameters such as the the shape of the presynaptic place fields and the animal’s velocity. We show explicitly that the place fields become broader if the animal has a higher velocity, and that LTD far away from the instructive signal has a slow convergence time to the fixed point. The resulting place fields also depend on the parameters of the eligibility traces and the instructive signal, parameters that may be inferred directly by experiments.

## 2 Model

### 2.1 Setup

The general framework for the model is as follows, emulating the setup for recent experiments in hippocampus [17, 18]. A mouse runs along a treadmill of length L at velocity *v*. Experimenters record from a postsynaptic CA1 place cell which receives inputs from N presynaptic CA3 inputs. As the animal runs along the track, a dendritic calcium spike (”plateau potential”) is artificially triggered in the CA1 cell, at the same location each lap, in order to induce learning. The CA3 inputs are themselves place fields which tile the length L of the running track (Figure 1). The firing rate *R_i_* of each CA3 input *i* is modeled as a Gaussian function of position:

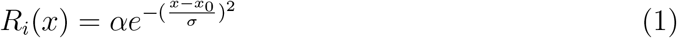

**Figure 1:**
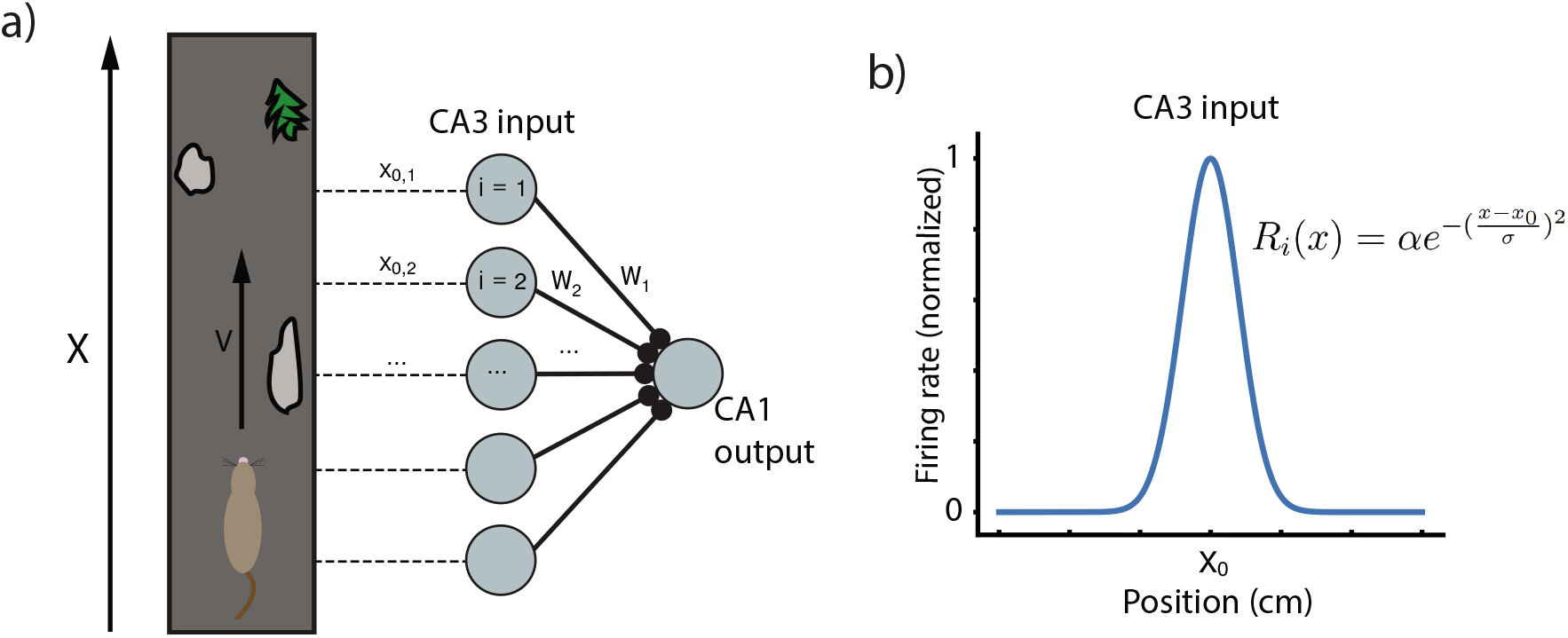
Running track and model network. a) A mouse runs at velocity v along a a running track with locations marked by unique features. Inside the mouse hippocampus, N CA3 place cells have activity peaks at different locations along the track, and synapse onto a single postsynaptic CA1 cell. b) The CA3 place cells considered here are modeled as simple Gaussians centered at evenly spaced locations along the running track.

Where *x*_0_ is the center of the given input receptive field. The CA1 output has a ramp potential (i.e. the membrane potential relative to rest, low-pass filtered to eliminate spikes, see [18]) determined by the sum of its synaptic input:

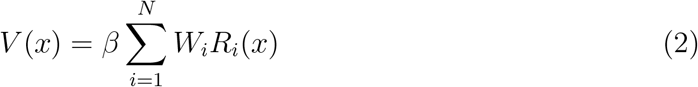

Each synapse in our model produces two traces, one for LTP and one for LTD, upon presynaptic firing *R_i_*. The equation for each trace 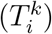 has the form:

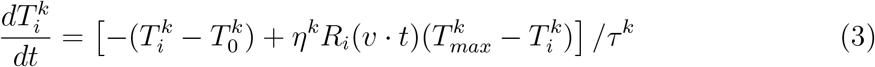

where 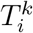 is a trace for synapse *i, k* indicates either LTP or LTD, 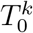 is the basal value of the trace for that synapse, 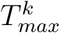 is the maximal value of the trace, *τ^k^* the time constant, and *η^k^* and activation rate constant. By a simple change of variables *T_i_* → (*T_i_* + *T*_0_), and *T_max_* → *T_max_* + *T*_0_, one gets the slightly simpler equation:

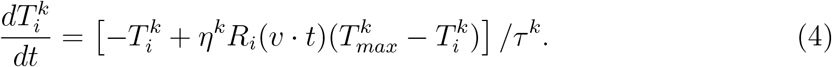

These traces act as transient markers of the presynaptic firing history, allowing the network to bridge events that occur within the temporal scale of the trace (*τ^k^*). The ODE dictating the traces has two terms, the first of which is a decay term - in the absence of presynaptic firing, this term causes the traces return to their basal level at a rate determined by the time constant *τ^k^*. The second term is an activation term, wherein presynaptic firing causes the traces to approach their saturation value 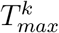. The shape of the trace depends on the trace parameters and on the shape and location of the place field of the presynaptic neuron to synapse *i*. Some examples of such traces can be found in Figure 2b.

**Figure 2:**
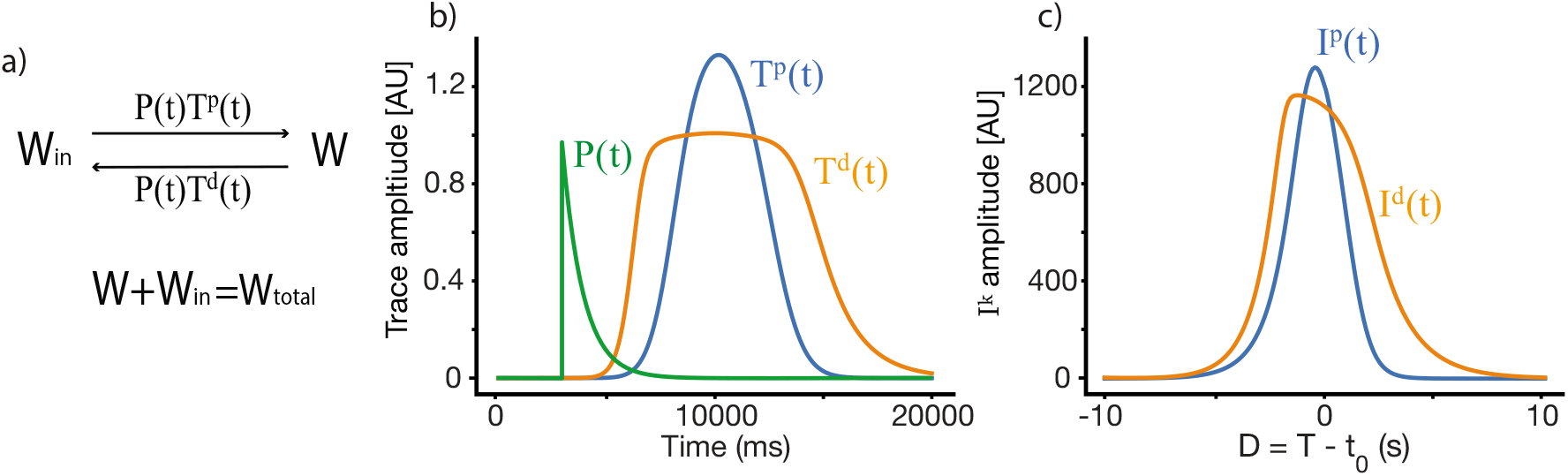
Weight dynamics. a) A simple dynamical model of synaptic plasticity, where both LTP and LTD require an overlap between a trace variable *T^k^* and an instructive signal *P. W* is the active synaptic weight, and *W_in_* are internalized resources, the total synaptic weight is conserved. b) An illustration of the synaptic plasticity traces and the instructive signal. c) The overlap *I^k^* between the traces and the instructive signal, iterated over locations of the instructive signal, where *D* = *T* − *t*_0_ is the displacement between the start of the instructive signal and the center of the presynaptic place field in units of time.

For a specific functional shape of an input place field *R*(*x*), the place field in time has the form *R*(*v* · *t* − *x*_0_) = *R*(*v* · (*t* − *t*_0_)) where *x*_0_ indicates the place field center in space. Therefore traces too can be written as 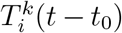. The traces interact with an ”instructive signal” P(t) which is triggered by the induced plateau potential:

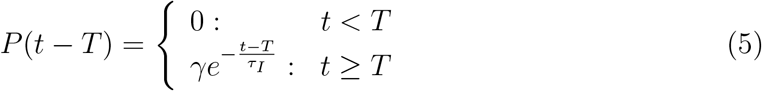

Where *T* marks the time of induction of the plateau potential. This instructive signal is global, acting across all synapses in tandem with the synapse specific traces.

Our learning rule is a very simple induction model which depends on above described synaptic traces 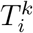, instructive signal *P*(*t*), and assumes a conserved resource that can become synaptic efficacy (see Figure 2a). These assumptions produce a simple ODE for the dynamics of synaptic plasticity:

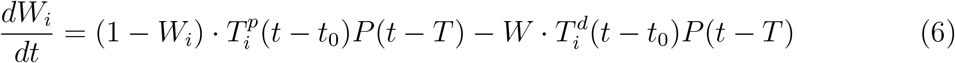

Where again we denote the start time of the instructive signal as *T* and the temporal center of the presynaptic place field as *t*_0_. By changing variables such that *t* → *t* + *t*_0_ we get:

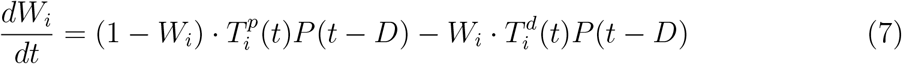

where *D* = *T* − *t*_0_ is the displacement between the start of the instructive signal and the center of the presynaptic place field in units of time. These presynaptic place fields are tiled along the length L of the running track, therefore for each synapse *i, D* is different, but simply linearly shifted. If the set of presynaptic neurons have centers that are equally and linearly spaced with a spacing Δ*x*, starting at *x*_0_ then *t*_0_(*i*) = *T* − (*x*_0_(*i*) + *i* · Δ*x*)/*v*), where *t*_0_(*i*) is the center in time or the presynaptic receptive field of synapse *i*. Similarly one could write *D* = *D*(*i*), where each version of D has he same order but is simply shifted.

### 2.2 General solution and fixed point

To find the fixed point solution of our learning rule, we can integrate over a single trial and assume that during that trial, *W* does not change significantly. The dynamics will therefore depend on the integral of the overlap between the traces and the instructive signal (Figure 2c):

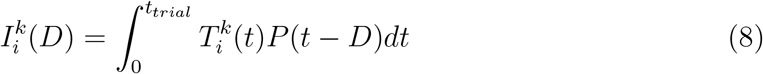

where *k* ∈ (*p, d*). The fixed point of equation 6 is therefore:

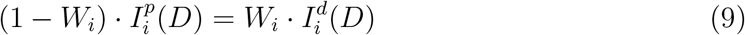

This implies that the fixed point of *W_i_* is simply:

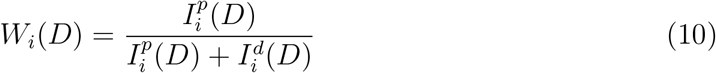

Practically speaking, this fixed point *W_i_*(*D*) gives us the final weights of all the synapses in response to an instructive signal presented at time T as a function of their temporal distance to this instructive signal, *D* = *T* − *t*_0_. This fixed point can be calculated numerically in the general case, but can also be calculated analytically under certain conditions (see Results and Methods). The fixed point can also be described in terms of spatial dependence, the units for which can be obtained in the case of constant velocity simply by multiplying *D* by the animals velocity *v*.

Equation 10 implies that with linear induction of traces, as assumed in this section, the parameters of the two traces must be different if we want the weights to depend on the displacement between the place field center and the instructive signal. If the traces are identical for all *D, W_i_*(*D*) = 0.5 for all *D*. If they have the same functional form, but a different scale such that *I^p^* = *k* · *I^d^* then *W_i_*(*D*) = *k*/(1 + *k*) for every *D*. For the formulation of traces in equation 3, this would occur, for example, if the parameters *η^d^* = *η^p^* and *τ_d_* = *τ_p_* but 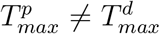. In order for the weights to produce place fields that are selective for a certain range of D, the overlap *I^d^* should generally be broader and shallower than the overlap *I^p^*. This can be implemented directly by choosing appropriately different trace parameters for LTD and LTP, and/or by including a basal level of LTD 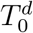.

## 3 Results

### 3.1 Linear Track

To investigate the evolution of the weights and their convergence to fixed points, we first consider the case of a linear track. For a linear track, we assume after each lap the animal ”restarts”, such that for a single trial, previous traversals of the track do not interfere. By calculating equation 10 for a set input receptive field shape *R* and a set instructive signal *P*, we can numerically solve for the fixed points. The resulting steady state place field shapes depend on the different parameters of the LTP and LTD traces. In Figure 3 we show different place field fixed points, with the same presynaptic place field and the same velocity, but with different trace parameters, as indicated above each subplot. One can observe that the width, selectivity, symmetry and overall shape of the place fields significantly depends on these parameters. From experiments we know that the shape of the fixed point should be such that it is maximal near *D* = 0 and gets smaller, approximately symmetrically as *D* becomes larger than zero. For a large enough *D, W_i_* should be close to zero. Given these experimental observations, one can estimate the different trace and instructive signal parameters that are consistent with experiments.

**Figure 3:**
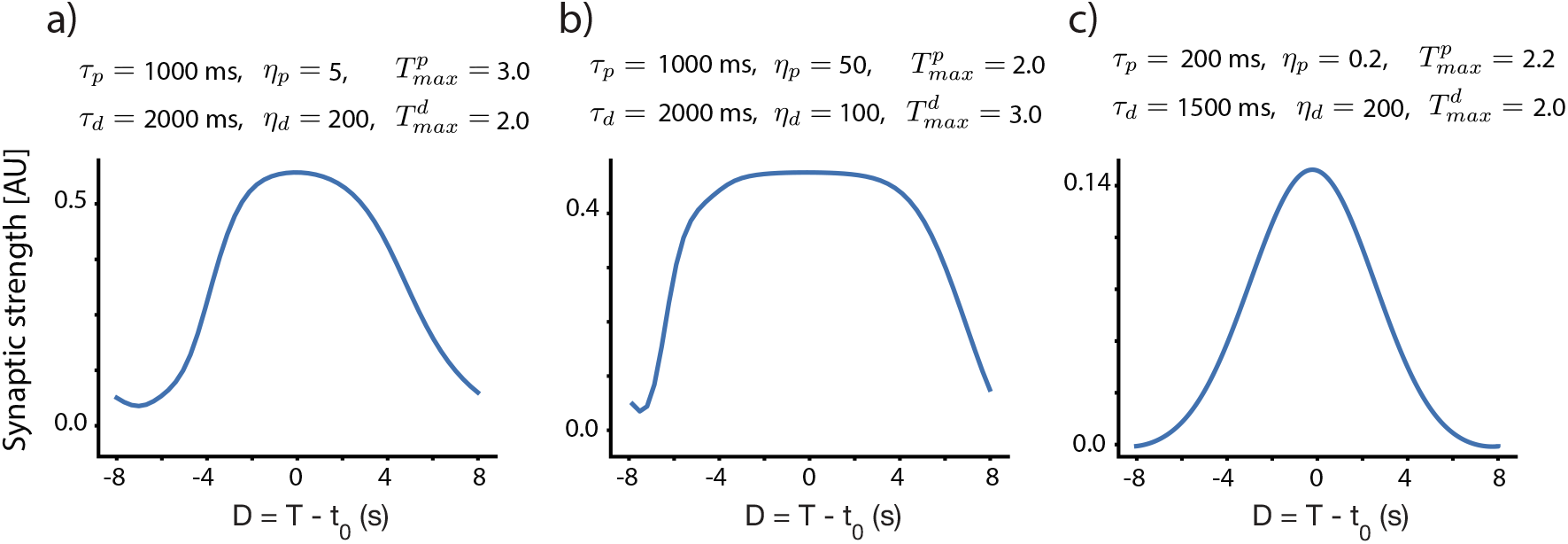
Fixed point place field structure. a-c) Fixed point weights as a function of D for different sets of parameters. The velocity in all these subplots is identical (v = 0.116 m/s), and the same presynaptic Gaussian place field is used, with a SD width of 0.21 m. All other parameters are indicated above each subplot.

These numerical calculations can then be checked by performing simulations where *W_i_* is explicitly updated at every time-step using equation 7. For the following simulations, we use the same parameters as in Figure 3c. We examine here two different cases. In the first case, initial weights are set to zero, and so the weights smoothly converge to their unimodal fixed point as the mouse repeats laps along the track. Figure 4a shows the fixed point CA1 output voltage (equation 2) that results from this case.

**Figure 4:**
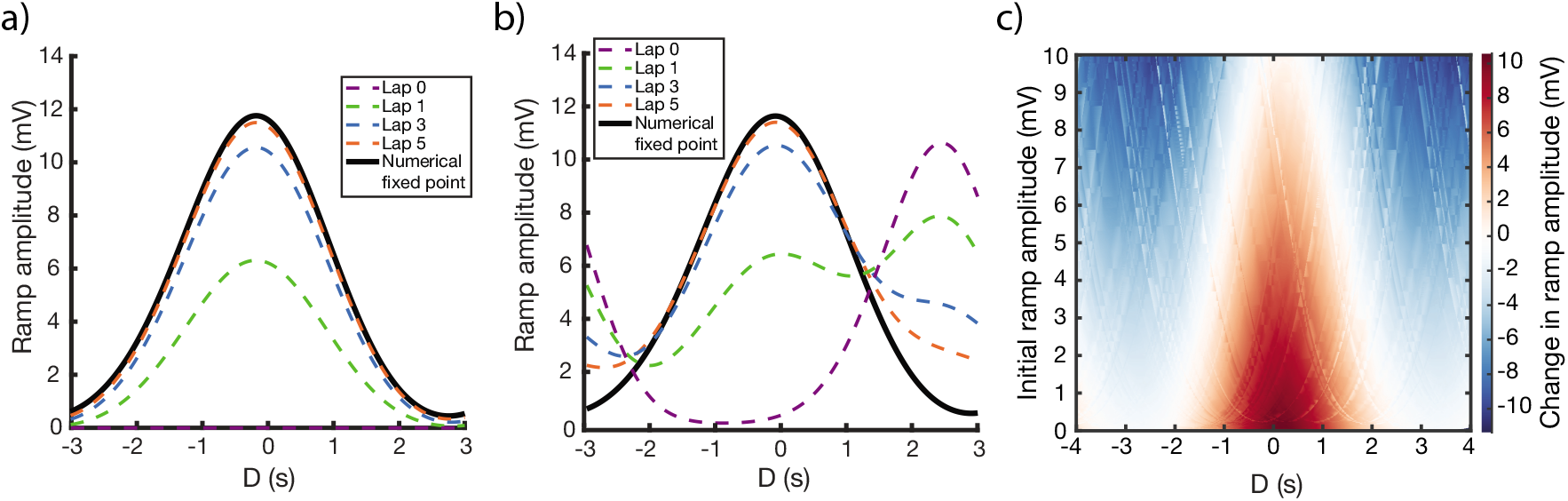
Linear track simulations. a,b) Simulated evolution of weights over laps (dashed lines) and the numerically calculated fixed point weights (solid lines) as a function of D. For a), initial weights are set to zero. For b), weights are initialized such that there is a preexisting place field at D = 3.5s. c) Change in ramp amplitude as a function of D and the initial ramp amplitude, calculated by iterating over 100 random selections of initial/new place field locations, and performing interpolation on the results.

In the second case (Figure 4b), the weights are initialized such that there is a preexisting place field in the CA1 output. Over the course of learning, interaction between the traces and the instructive signal cause the new place field (centered at the location of the instructive signal, D = 0) to potentiate and the preexisting place field to depress. The further away the initial place field is from the new place field, the longer it will take the initial place field to depress, owing to a smaller overlap (*I_k_*) between the traces and the instructive signal. However, the fixed point of the weights are irrespective of the initial place field, so given enough laps, the initial place field will always disappear, and the solution will become unimodal.

Note that in both of these cases, and despite the assumptions made in deriving the fixed point solution of equation 10, the numerical and simulation results are nearly identical.

Running multiple simulations while iterating over initial place field locations and instructive signal locations, one can plot the change in place field ramp amplitude as a function of initial place field ramp amplitude and the time relative to plateau onset. This creates (Figure 4c) a generalized response curve that predicts the results of inducing a plateau potential, in a given location and with a given initial ramp voltage. This generalized curve can be modified by changing parameters of the model.

### 3.2 Circular track

A similar exercise can be performed assuming a circular track with continuous running, where the instructive signal/traces can span across laps and place fields are periodic. We can think of each cycle as another iteration in which the final value of traces on the previous run is their initial value of the current run. In this case it is more complicated to calculate the numerical fixed point, but one way to do so is to consider an infinite number of iterations, and use the convergence of the infinite series (see subsection 4.2).

In order to illustrate this graphically, we show in Figure 5a the traces for a lap >> 1, where due to the periodicity of the track, the traces’ (and the instructive signal’s) values at the end of the track are equal to those at the beginning of the track. This periodicity is also reflected in the overlaps *I_k_* (Figure 5b) which are used to calculate the fixed point. It turns out that for a wide range of parameters the circular track solution closely matches the linear track solution (Figure 5c). In particular, if the time constant of the traces is shorter than the remaining length of the track, the traces will decay before the track ends, meaning that the traces will be at or near zero at the start of the next lap in both the linear and circular case. As a result, the biggest differences between the circular and linear track fixed points occur near the start/end of the track, and when the trace time constants are long relative to the traversal time of the track. Other parameters show a larger difference between the circular and the linear environments. Different input place field shapes, such as rectangular, can also cause a separation in the linear and circular fixed point solutions because of the effects they have on the shape of the traces.

**Figure 5:**
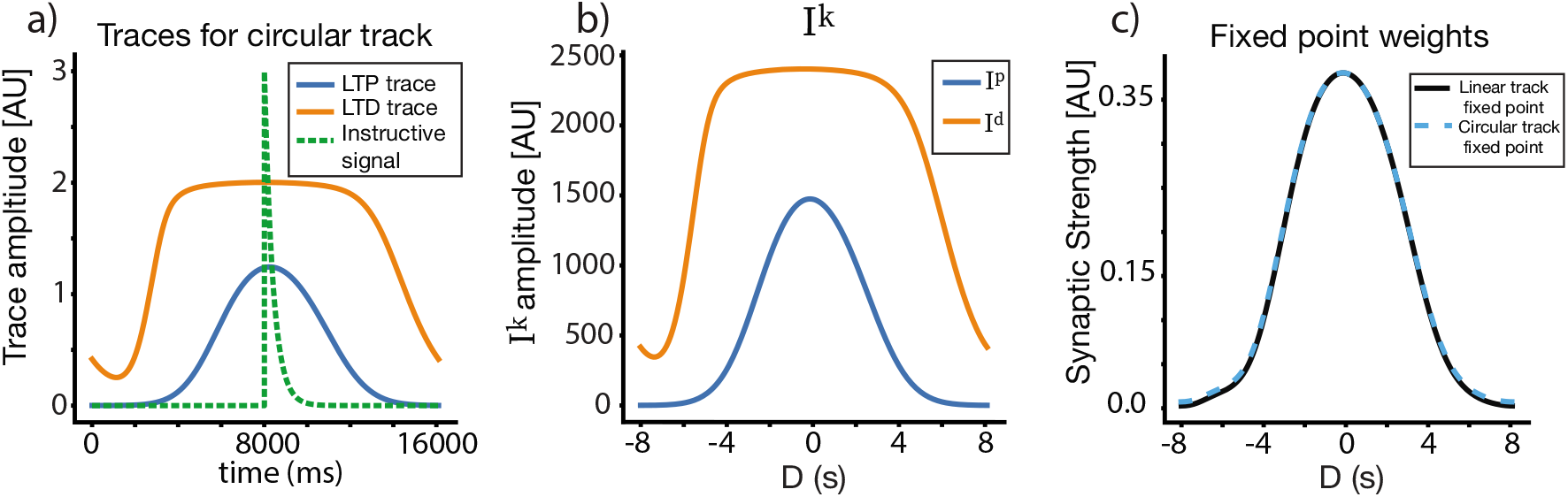
Place field plasticity on a circular track. a) Traces and instructive signals for a lap >> 1 on the circular track, as a function of time. b) *I_p_* and *I_d_* on circular track. c) Fixed point for weights on the circular track, compared to fixed point weights for the linear track. Notice that while the two solutions are nearly identical around *D* = 0, small deviations can be observed along the flanks near |*D*| = 8s.

### 3.3 Non-Linear trace activation

A further modification can be made to the circular track model by assuming a non-linear response of the traces to presynaptic firing. Here we assume that instead of the presynaptic firing rate *R*, directly activating the traces, this activation is non-linearly filtered though a function *F^k^*(*R*) where the index *k* can take the values *p* or *d* indicating a possibly different non-linearity for LTP and LTD traces. Thus the trace activation equation now takes the form:

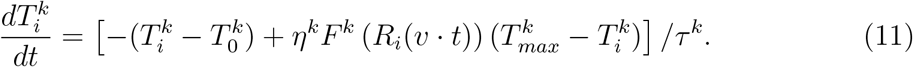

For simplicity we have chosen a thresholded linear form: *F^k^*(*x*) = *C^k^*[*x* − *θ^k^*]_+_. Practically it implies that the LTP and LTD traces see different effective presynaptic receptive fields, and aside from that, all the previous analysis still holds. The assumption of non-linear activation does not necessarily produce superior solutions (i.e. fixed points which more closely match experiment), but it allows for a different way of obtaining similar results. Examples of the solutions with non-linear traces on a circular track are shown on Figure 6. In figure 6a we show the effective receptive fields for the LTP trace (dashed red) and LTD trace (black) respectively. Apart from this difference, in this example, the parameters of the LTP and LTD traces are identical. The narrower LTP effective place field generates a narrower LTP trace, which allows for a selective receptive field (Fig 6c).

**Figure 6:**
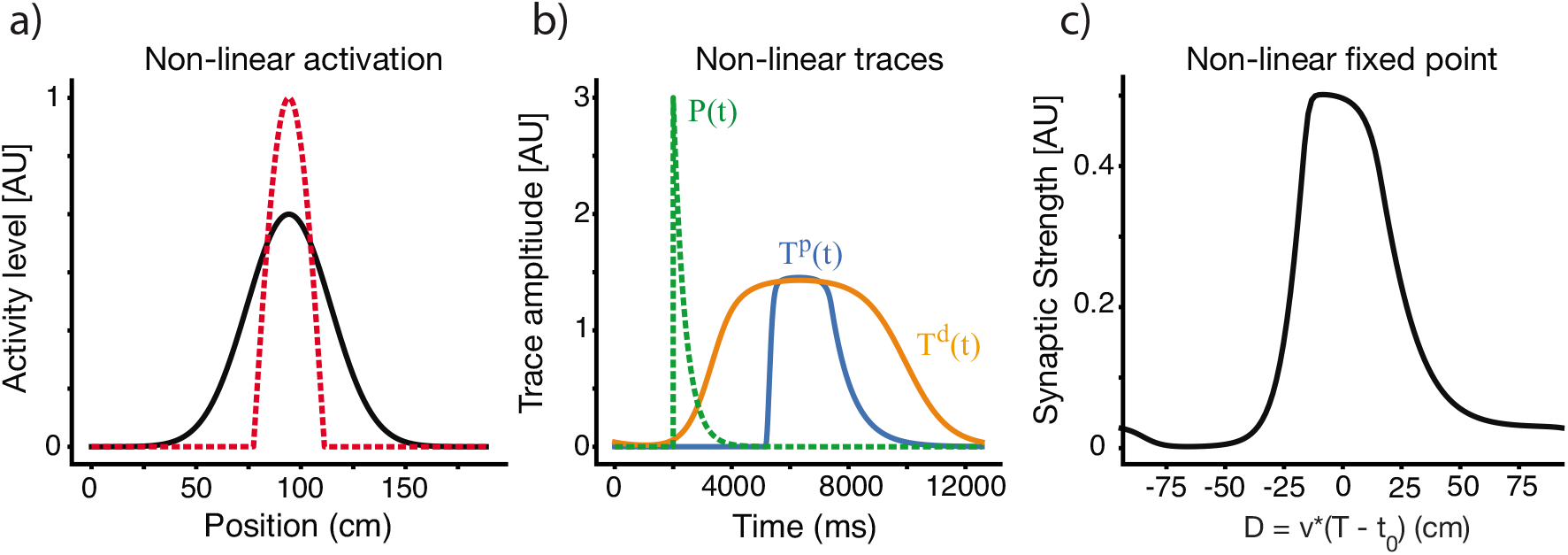
Place fields with non-linear traces. a) The effective presynaptic activity profiles that drive LTP and LTD traces are different. Here the LTD traces are driven by the linear RF as above (black) but LTP traces are driven by a non-linear modification of the RF (dashed-red). b) LTP (blue) and LTD (orange) traces in the non-linear case. The instructive signal in dashed green. c) The resulting fixed point of the weight vector. Results here are for *V* = 0.15 *m/s*.

### 3.4 Velocity Dependence

Until now, we have only shown results for one fixed running speed, but the shape of the resulting place fields depends on the velocity of the animal on the track. Figure 7 shows how the shape and selectivity of place fields depends on velocity for a given set of parameters. These results are consistent with experimental results, which show that as the velocity increases, the width increases and the selectivity of the resulting place field decreases. Qualitatively similar results are obtained for other parameter sets, and for non-linear traces. Velocity dependence is similar on the linear and circular tracks, so we only demonstrate it here for the circular track. Note that the fixed point of W here is plotted against D in terms of distance in centimeters, so that the X axis stays the same at different velocities. While at every velocity there is a single fixed point curve, experimentally we might not reach the fixed points because the convergence time might be large.

**Figure 7:**
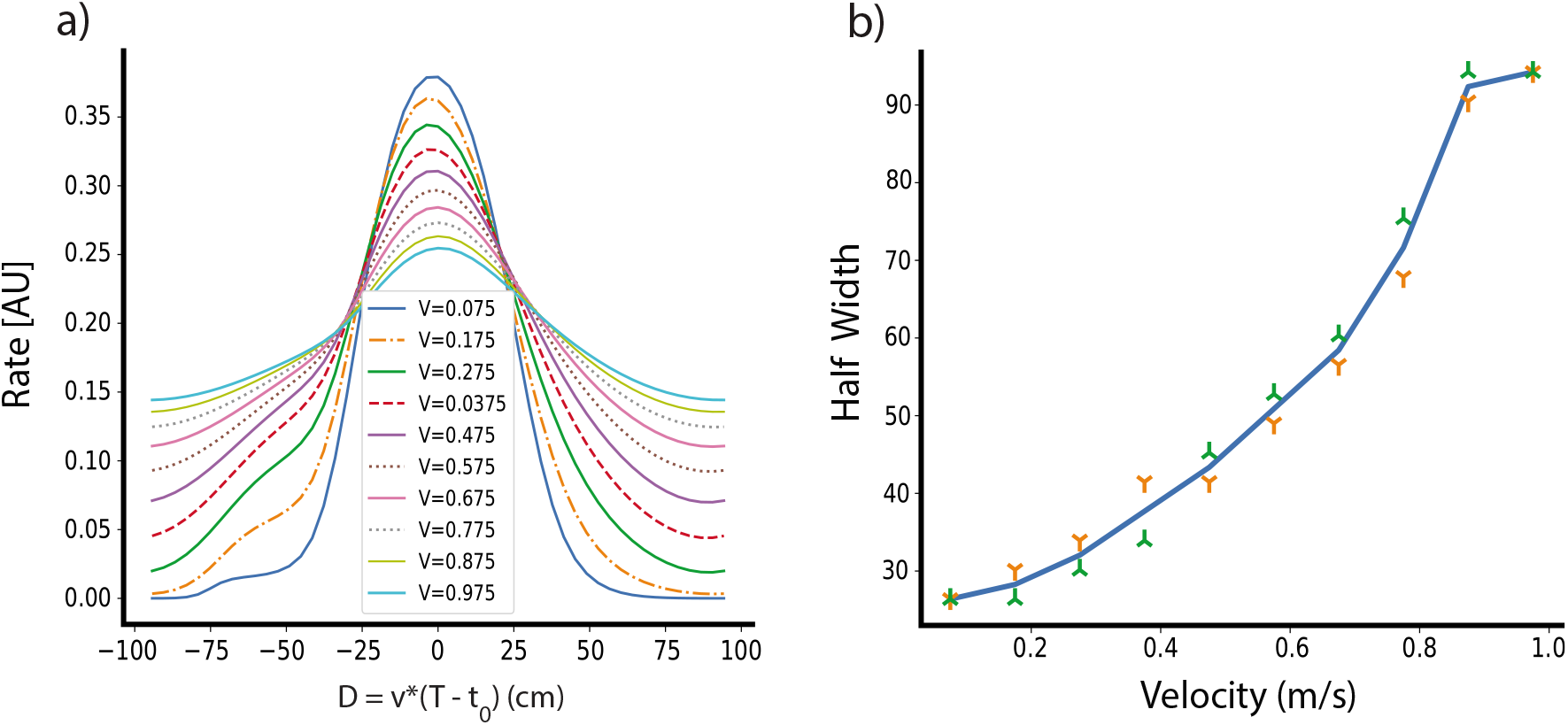
The dependence of place field shape on velocity. a) Fixed points of the weight vector as a function of D for different movement velocity. b) The dependence of the half width of the weights a fixed-point on the travel velocity. The green and orange symbols are for left and right widths, respectively.

### 3.5 Convergence Time

The fixed point of the learning rule produces a single place field centered at the location of the induced plateau potential, regardless of pre-existing place fields before learning. However, the rate at which learning converges depends on D - the larger absolute value of D, the longer the convergence time. This effect is present for both the potentiation of new place fields and the depression of old place fields. Figure 8a shows the simulated convergence of the weights towards their fixed points as a function of trial number and D, given zero initial weights. To predict the convergence time analytically, we use the approximation that W does not change significantly during a single trial to rewrite equation 6 as:

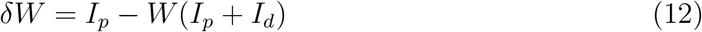

where *I_p_* and *I_d_* are defined as in equation 8. The solution to this difference equation is an exponential with the time constant 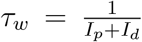, which gives us an estimate of the relative convergence time. Simulations show that the analytically predicted convergence time is closely related to the number of trials it actually takes for the simulated weights to reach 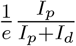 (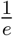, or approximately 36 percent of their fixed point values (Figure 8b) when starting from zero initial weights. Notably, the convergence time steeply rises for large absolute D, such that it takes around 3-5 times as many trials to converge for *D* = 8 s as it does for *D* = 0 s. For even larger D, *τ_w_* can become prohibitively long, meaning that the output can effectively maintain two place fields provided they are far enough apart from each other, as the old place field will depress incredibly slowly.

**Figure 8:**
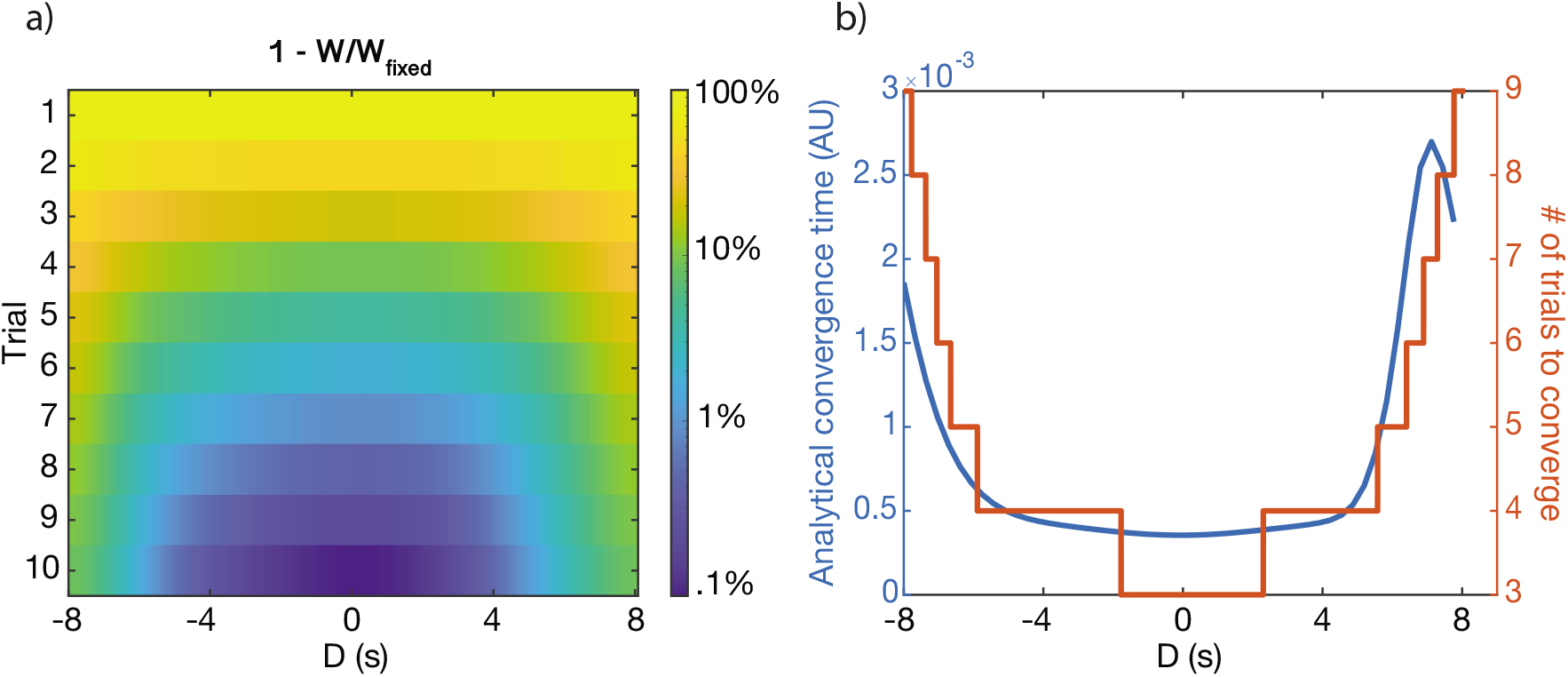
Convergence to solutions. a) The relative distance to the fixed point ((*W_fixed_* − *W*)/*W_fixed_*) as a function of *D*, shown as logarithmic heatmap. b) The convergence time, *τ_w_* (blue), and the number of simulated trials to reach *W_fixed_/e* (orange) as a function of *D* (in seconds). Notably, in both cases the convergence time rises steeply beyond a certain value of D. Both cases start from the initial condition *W_i_* = 0.

### 3.6 Analytical solution for rectangular place fields

The numerical integrals needed to find the traces 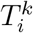, and their overlaps with the instructive signals, 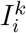 can in some cases be solved analytically. A simple example that can be solved analytically is when presynaptic receptive fields are rectangular. To solve for the fixed points analytically, we assume that input place fields are rectangular, starting at *t*_1_ and ending at *t*_2_, with an amplitude of *α*. In terms of the position variable these start at *vt*_1_ and end at *vt*_2_. Formally the place field has the form:

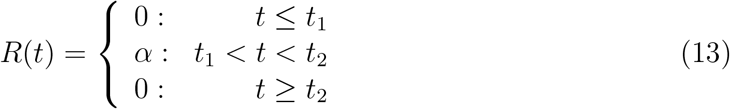

Using this simple form of *R*(*t*), we can explicitly solve 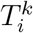 and therefore 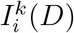 (8), in the case of the linear track. By performing these calculations (see Methods), we arrive at the fixed point 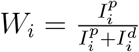. The final analytical form of the fixed point depends on the activation rate (*η*), saturation values 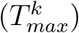 and time constants (*τ^k^*) of the traces, the location (*t*_1_ and *t*_2_) and amplitude (*α*) of the receptive field, location (*t*_0_) and time constant (*τ_p_*) of the instructive signal, and on the duration of the lap *t_trial_*. We also include in this analysis a basal level of LTD 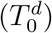, which smooths over some of the artifacts of the rectangular input place field, and makes the solution more similar to the solutions with Gaussian input place fields demonstrated above. Figure 9 shows the results of the analytical solution. The fixed points found analytically match those found via simulations (and those found via numerical methods), given the same input PF and same parameters are used in both cases (Figure 9b).

**Figure 9:**
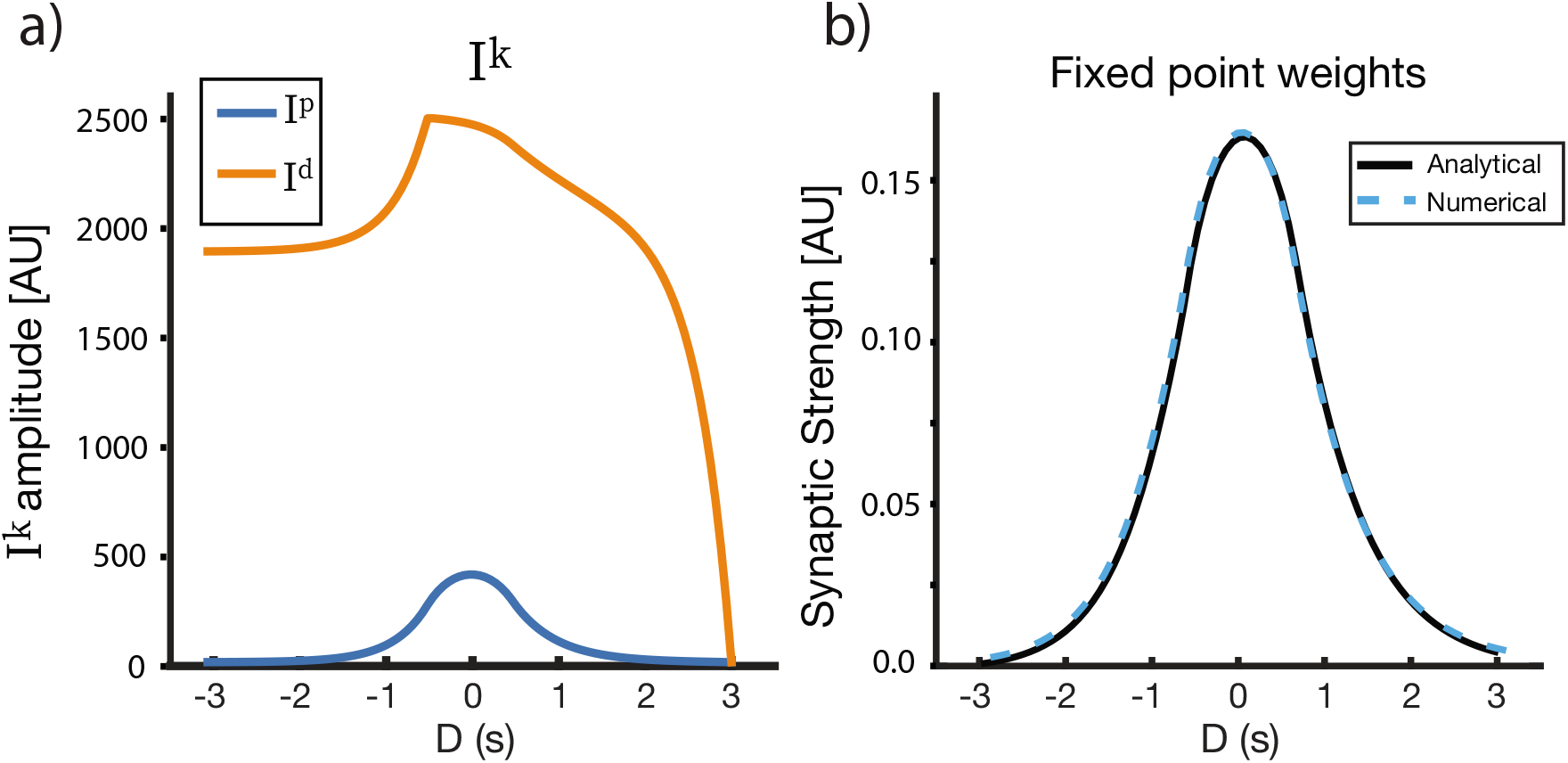
Analytical solutions for a rectangular receptive field. a) *I_p_, I_d_*, and b) *W_fixed_* as a function of D. Here, the LTD trace has a basal level 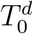, which creates an additional overlap 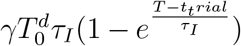 (see Methods). The resulting fixed point is close to symmetric around D = 0, and its properties can be modified via the model parameters. The parameters used for the figure are as follows: *τ_p_* = 500ms, *τ_d_* = 1500ms, *η_p_* = .25, *η_d_* = 200, 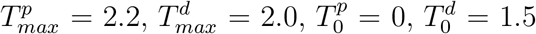.

## 4 Methods

### 4.1 General trace solution

We can find a closed form solution for the traces by directly integrating equation 3. Equation 3 is a non-homogeneous linear first order ODE, which can be solved using an integration factor *U*(*t*). Where:

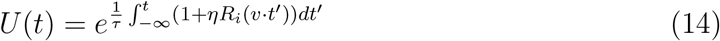

The solution for equation 3 then has the form:

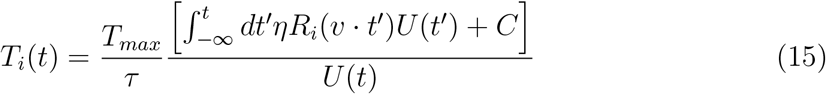

Given *R_i_*(*v* · *t′*), one can use equation 15 to solve for *T_i_*(*t*), and therefore 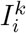, and therefore *W_i_*. Some choices of *R_i_*(*v* · *t′*) are more analytically tractable than others, hence the choice of rectangular presynaptic place fields for analysis (equation 13), as they allow for a closed form solution to this equation.

### 4.2 Circular track

For a circular track, we must account for the wraparound of both the traces and the instructive signal in our calculations. If we define as *t_max_* as the time it takes to do one cycle, then the total time up to time *t* can be rewritten as *t* = *n* · *t_max_ + q*, where *n* is the number of fully completed cycles. On the *n* + 1 run the initial condition denoted by *C_n_* is *T_i_*(*n* · *t_max_*). Using these definitions, and the periodicity of the place cell with a period *t_max_*, the solution in equation 15 is:

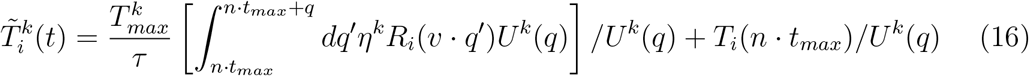

where the index *k* is *p* or *d* for LTP and LTD traces, respectively. The first part on the right is the same for *q* in each period, and independent of *T_max_*. The second part iteratively forms a series such that: 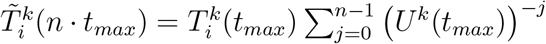 which in the infinite limit converges to: *T_i_*(*t_max_*)/(1 − *U*(*t_max_*)^−1^) This implies that the solution for the circular track converges to:

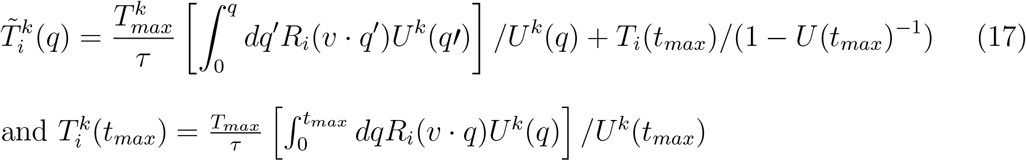

### 4.3 Convergence time

Explicitly, the solution to equation 12 is:

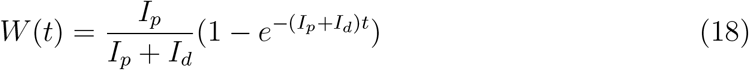

Though our assumption to reach this equation (that W does not change much from trial to trial) is not necessarily true for all D, it is close enough to make this a good first approximation of W(t) (and therefore 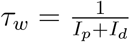 a good approximation of a convergence time constant). For |*D*| >> 0, the weights do indeed change quite slowly, and equation 18 is a reasonable estimate of the weight dynamics.

### 4.4 Analytical solution for rectangular place fields

In this section we drop for simplicity the superscript for type of trace and the subscript for synapse index. Using the trace equation (3) and the above choice of *R*, the integration factor *U* (see section 4.1) has the form:

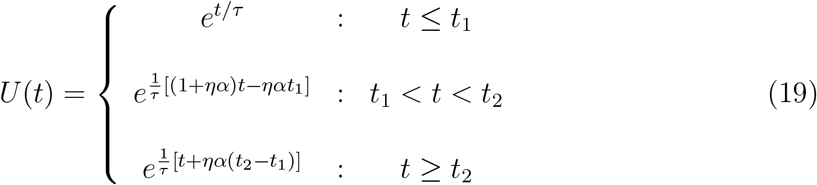

Define 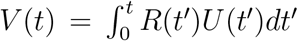. The solution (see equation 15) with *C* = 0 is simply *T*(*t*) = *ηT_max_V*(*t*)/(*U*(*t*)*τ*). Let’s now calculate *V*(*t*). For *t* ≤ *t*_1_, *V*(*t*) = 0, because *R*(*t*) is zero over this range. For *t*_1_ < *t* < *t*_2_:

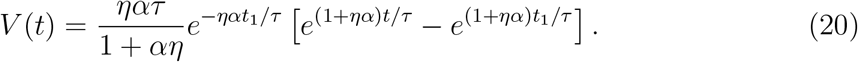

For *t > t*_2_: *V*(*t*) = *V*(*t*_2_)

Therefore:

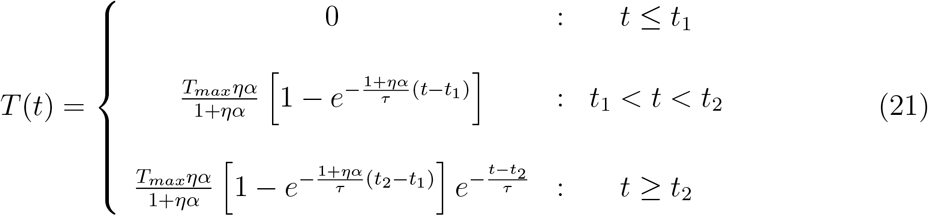

Recall that one may take the standard trace equation (equation 4) and transform it into the one with a basal level (equation 3) via the changes of variables *T_i_* → (*T_i_* − *T*_0_) and *T_max_* → *T_max_* − *T*_0_. So if we are to include a basal level in our analytical solution, all the *T_max_* in equation 21 become *T_max_* − *T*_0_, and the constant *T*_0_ is added to the right side of the equation in all three cases.

Now that *T*(*t*) has been solved, one may find the fixed point of *W_i_* (equation 10) by solving for 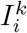 (equation 8). Assume that we have an instructive signal of the form:

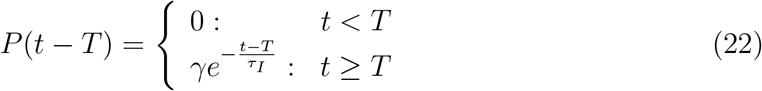

Where *T* is the time of the start of the instructive signal. Since 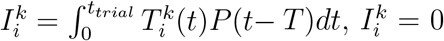 for both 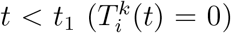 and *t* < *T* (*P*(*t* − *T*) = 0). This leaves us with three cases to integrate over: *T < t*_1_, *t*_1_ ≤ *T* < *t*_2_, and *t*_2_ ≤ *T < t_trial_*.

For *T < t*_1_:

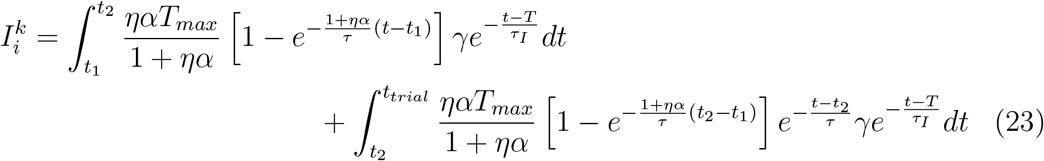

which can alternatively be written as:

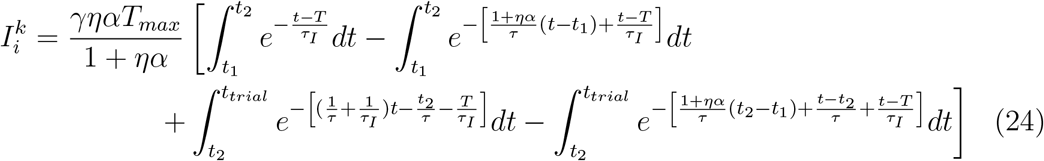

The two other cases (*t*_1_ ≤ *T < t*_2_ and *t*_2_ ≤ *T < t_trial_*) can be solved via the equation above by substituting the limits of integration as appropriate. Following this prescription leads to the following solutions:

For *T < t*_1_:

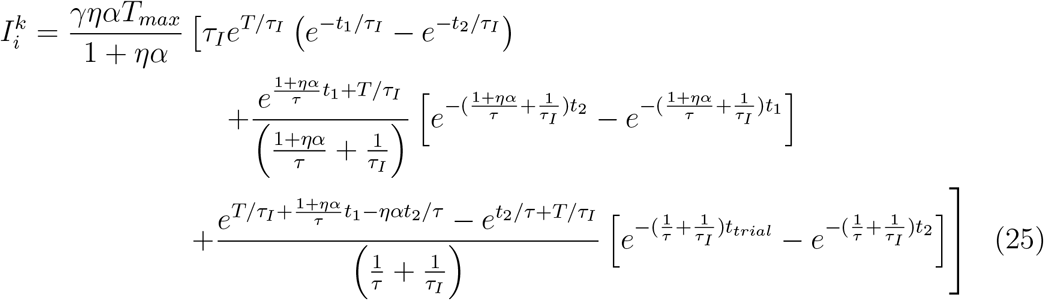

For *t*_1_ ≤ *T < t*_2_:

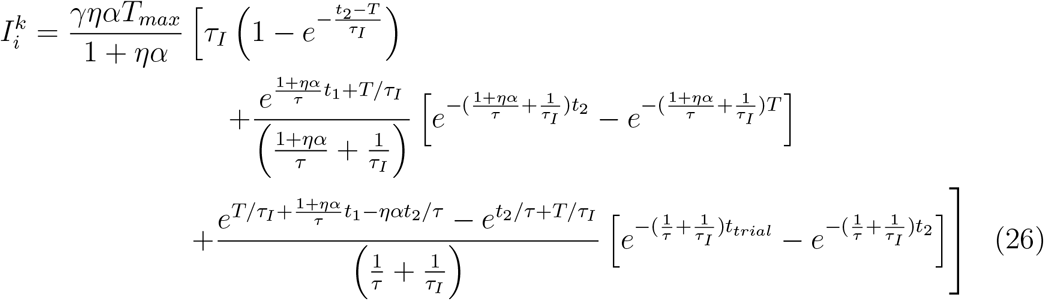

For *t*_2_ ≤ *T < t_trial_*:

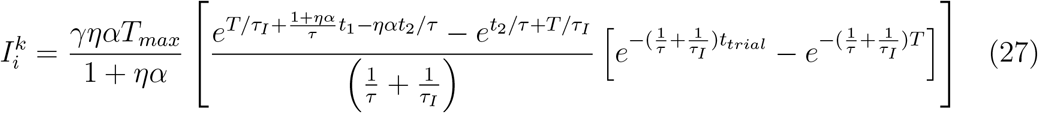

For traces with a basal level, each of three cases has an additional basal overlap 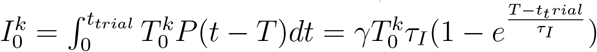.

The fixed point can then be calculated by plugging in these expressions for 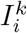 into 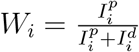 (equation 10), for each of the three cases. Figure 9 shows the fixed point solution that results from these calculations, assuming a non-zero basal level for the LTD trace.

## 5 Discussion

Synaptic plasticity that operates on behavioral time scales has been shown to determine place field plasticity in CA1 neurons. The underlying structure of such plasticity is significantly different than the commonly studied Hebbian forms of plasticity that assume near temporal coincidence of pre and postsynaptic activity. Experimental results in CA1 using both in vivo and slice preparations strongly suggest that presynaptic activity generates synaptic eligibility traces for both LTP and LTD [17, 19]. Prior modeling work [18] shows, using simulations, that these two opposing traces, combined with a weight dependence can account for the place field plasticity observed in vivo.

Here we proposed a simple formulation for two-factor plasticity [20] that depends on eligibility traces for both LTP and LTD, an instructive signal and weight dependence. We have shown that this rule can be mathematically analyzed to yield a simple expression for the fixed points of these place fields. These results show that place fields can have spatial selectivity only if the traces have essentially different temporal dynamics, or if they are induced non-linearly, and with different non-linearities. Such results are general, beyond the specific shapes of presynaptic place fields and the model’s parameters. The general rule in both cases is that for obtaining selective place fields, the overlap between the LTD trace and the instructive signal should have a broader shape the the overlap between the LTP trace and the instructive signal; the more pronounced this difference, the sharper the place field. By assuming specific shapes of presynaptic activity and of the instructive signal, we can calculate exactly the shape of the learned postsynaptic place fields at steady state and can estimate the local convergence rate to these shapes. We also used simulation to validate the analytical results and show that the convergence of place fields is rapid. In the case of a rectangular presynaptic place field a fully analytical solution is obtained.

The shape of the postsynaptic place field at steady state depends on many of the system parameters and assumptions, such as the shape of the presynaptic place field and the parameters that determine the dynamics of the traces or the instructive signal. One additional parameter that significantly affects the shape of the place field is the animal’s velocity. If the animal moves through space slowly, place fields are more local and selective, while a fast velocity results in non-local and broad place fields. These results generalize beyond a narrow set of parameters, and can therefore be used to experimentally validate the model.

We have also shown that the weakening of prior place fields in old locations is significantly slower than the strengthening of the previously weak efficacies in the new location. The determining factor for this rate of change is the overlap between both eligibility traces and the instructive signal. Locations near the peak of the instructive signal are the ones that get enhanced, as in such locations there is a maximal overlap between the traces and the instructive signal, and therefore a fast convergence rate. On the other hand, locations which might have strong initial weights but are far from the instructive signal get weakened. For such signals the overlap between the instructive signal and both traces is much smaller and therefore the convergence to the fixed point is also much slower.

In this formulation we have assumed that the synaptic efficacy dynamics are weight dependent. Such weight dependence can arise from an assumption of a conserved quantity, for example a conserved number of receptors in the membrane and an internal synaptic store. This assumption is also motivated by a previous weight dependent model, that compares well with experimental results [18]. The existence of fixed points in our model, and their shape critically depends on this assumption.

In our solution we assume that the animal’s velocity is constant throughout, though in a real environment this is not the case. Animals change their running speed as they traverse the track, speeding up and slowing down and even sometimes stopping to eat. Such varying velocity implies that there is no true fixed point with this model, and that the place fields fluctuate around some mean fixed point. For a changing velocity, one can perform simulations using experimentally obtained velocity profiles instead of an analytical solutions [18]. Such results are more biologically realistic but are much more complex and offer less intuition. It is feasible that using the animal’s velocity statistics, the mean around which the place fields fluctuate, the variance with which they fluctuate, etc. can be estimated.

The inclusion of eligibility traces in our learning rule is essential, as it allows the model to associate activity with temporally distal instructive signals and thereby solve the temporal credit assignment problem. Traditional models of learning such as Hebbian plasticity or STDP are insufficient to describe plasticity which occurs on behavioral time scales, as they are restricted to learning tight temporal correlations of activity. Previous characterizations of eligibility traces have generally depended on both pre- and postsynaptic activity, where the conversion of traces to actual synaptic efficacies depended on a third factor such as a reward signal [7, 8, 12, 21, 13, 20]. In contrast, the model presented here is essentially a two-factor rule, in which traces depend only on presynaptic activity, and weight changes depend on both traces and the instructive signal. For feed forward circuits with a supervised or reinforcement type objective (such as the hippocampal circuit we describe in our model), there is no real need for three factor learning since the circuit tries to find an association between two external factors a stimulus and some form of instructive signal. Indeed, previous work has shown that experimentally observed behavioral timescale synaptic plasticity in CA1 is in fact inconsistent with three-factor rules which depend on synchronous pre- and postsynaptic activity [18]. The observation that place field plasticity in CA1 neurons can be described by a two-factor rule actually makes its analysis simpler, enabling the detailed results found here. In other systems, such as sensory systems (for which previous eligibility trace theories were developed), the aim is to alter the dynamics of a recurrent network, so for such models three-factor might indeed be necessary [12, 8, 13]. Regardless, both two- and three-factor eligibility trace models provide a simple and mathematically tractable explanation for learning that occurs on behavioral time scales.

## 6 Acknowledgments

We would like to thank Jeff Magee, Sandro Romani and Aaron Milstein for their inspirational work and the productive discussions we had regarding it. We would also like to thank the following agencies and grants for funding this work - NIH: Harel Z Shouval, R01 EB022891; ONR: Harel Z Shouval, N00014-16-1-2327.

